# SARS-CoV-2 Omicron infection augments the magnitude and durability of systemic and mucosal immunity in triple-dose CoronaVac recipients

**DOI:** 10.1101/2023.09.08.556870

**Authors:** Yuxin Chen, Tiantian Zhao, Lin Chen, Guozhi Jiang, Yu Geng, Wanting Li, Shengxia Yin, Xin Tong, Yue Tao, Jun Ni, Qiuhan Lu, Mingzhe Ning, Chao Wu

## Abstract

**Background:** The inactivated whole-virion vaccine, CoronaVac, is one of the most widely used coronavirus disease 2019 (COVID-19) vaccines worldwide. There is a paucity of data indicating the durability of the immune response and the impact of immune imprinting induced by CoronaVac upon Omicron breakthrough infection.

**Methods:** In this prospective cohort study, 41 recipients of triple-dose CoronaVac and 14 unvaccinated individuals were recruited. We comprehensively profiled adaptive immune parameters in both groups, including spike-specific immunoglobulin (Ig) G and IgA titers, neutralizing activity, B cells, follicular helper T (Tfh) cells, CD4^+^ and CD8^+^ T cells, and their memory subpopulations at 12 months after the third booster dose and at 4 weeks and 20 weeks after Omicron BA.5 infection.

**Results:** Twelve months after the third CoronaVac vaccination, spike-specific antibody and cellular responses were detectable in most vaccinated individuals. BA.5 infection significantly augmented the magnitude, cross-reactivity and durability of serum neutralization activities, Fc-mediated phagocytosis, and nasal spike-specific IgA responses, memory B cells, activated Tfh cells memory CD4+ T cells, and memory CD8+ T cells for both the ancestral strain and Omicron subvariants, compared to unvaccinated individuals. Notably, the increase in BA.5-specific immunity after breakthrough infection was consistently higher than for the ancestral strain, suggesting no evidence of immune imprinting. Immune landscape analyses showed vaccinated individuals have better synchronization of multiple immune components than unvaccinated individuals upon heterologous SARS-CoV-2 infection.

**Conclusion:** Our data provides detailed insight into the protective role of inactivated COVID-19 vaccine in shaping humoral and cellular immune responses to heterologous Omicron infection.

**Trial registration:** ClinicalTrials.gov NCT05680896

**Funding:** This study was supported by the National Natural Science Foundation of China (92269118, 92269205), Nanjing Important Science & Technology Specific Projects (2021-11005), Scientific Research Project of Jiangsu Health Commission (M2022013), Clinical Trials from the Affiliated Drum Tower Hospital, Medical School of Nanjing University (2021-LCYJ-PY-9), and Jiangsu graduate practice innovation project (JX22013929).

## Introduction

Three years after the coronavirus disease 2019 (COVID-19) outbreak caused by severe acute respiratory syndrome coronavirus 2 (SARS-CoV-2), remarkable collaborative efforts have been made to mitigate the pandemic. The worldwide COVID-19 vaccination campaign demonstrated outstanding effects on reducing the severity of SARS-CoV-2 infection.^1-3^ CoronaVac, a whole-virion inactivated vaccine produced by Sinovac, is the most widely offered COVID-19 vaccine globally. Although the diminished immunity and emergence of new viral variants have fueled an increase in the frequency of breakthrough infections, a real-world efficacy analysis in China showed that the protection efficacy of the triple-dose inactivated vaccine against BA.2 infection was 74% against pneumonia and 93% against severe COVID-19.^4^ The triple-dose inactivated vaccine during the Omicron wave demonstrated an effectiveness of 69.6% against severe COVID-19.^3^

Understanding the underlying immunological mechanism accounting for this vaccine-derived protection is not only crucial for next-generation COVID-19 vaccine optimization, but also have important implications for future pandemic preparedness. In a prospective vaccine cohort, we previously analyzed the longitudinal immune responses elicited by the triple-dose inactivated COVID-19 vaccine up to 2 months, including serum neutralizing antibody activity and B and T cell immune memory responses.^2,5-7^ Since the adjustment of the public health policy for containing COVID-19 in China in December 2022, the subsequent SARS-CoV-2 Omicron variant wave resulted in large-scale breakthrough infection. In this study, we aimed to address two equally important issues. First, we examined whether CoronaVac-induced immune memory could still be effectively recalled up to 12 months after the third booster dose. The Omicron variant is associated with a marked capability to evade humoral immunity resulting from either prior natural infection or vaccination^8,9^. There are rising concerns regarding the immune imprinting induced by ancestral strain-based vaccinations, which might compromise the antibody response to Omicron-based boosters. Therefore, we also attempted to investigate whether the immune responses elicited by triple-dose CoronaVac had any beneficial role in the anti-viral responses to subsequent Omicron infection.

In this study, we investigated these key questions by measuring SARS-CoV-2-specific dynamic immune trajectories, including antibody responses, Fc-mediated effector function, neutralization capacity, and cellular responses after Omicron BA.5 infection among recipients of triple-dose CoronaVac and unvaccinated individuals. Our data provide detailed insight into the favorable role of pre-existing immune responses elicited by triple-dose CoronaVac, which are capable of potently recalling SARS-CoV-2 immunity with cross-recognition to Omicron variants.

## Results

### Study cohort and study design

We recruited a total of 55 participants, including 41 recipients of triple-dose CoronaVac and 14 individuals who were not vaccinated against COVID-19 (**Figure 1a**). Omicron BA.5 infection was confirmed in all individuals by next-generation sequencing. The demographic and clinical characteristics of the cohort are summarized in **Figure 1b**.

**Figure 1.**
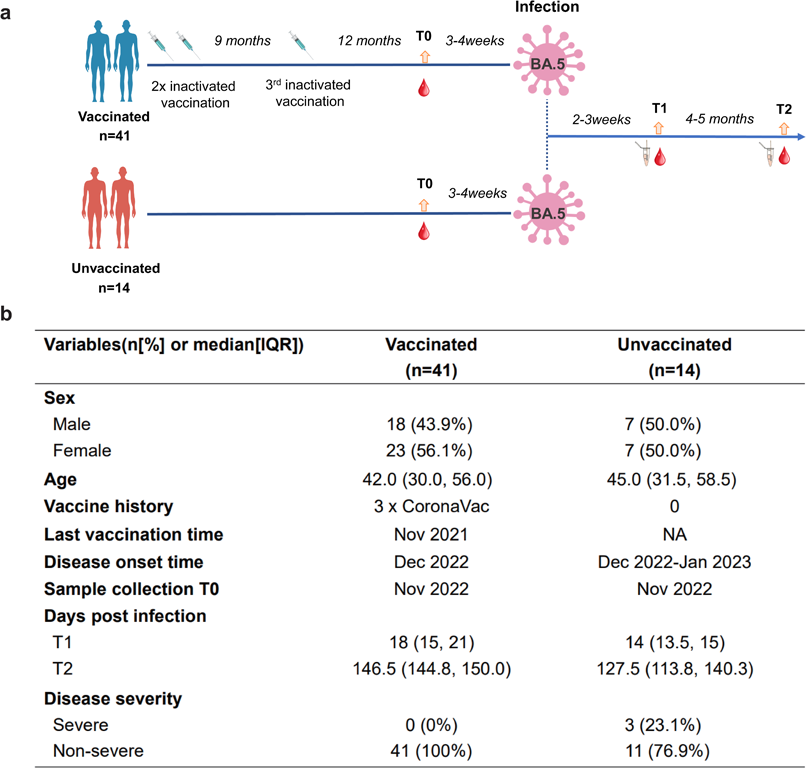
Study design and demographic characteristics of our cohort. (**a**) Study design of vaccinated individuals and unvaccinated individuals with Omicron BA.5 infection. Vaccination, infection and blood draw timeline were also noted. (**b**) The demographic and clinical characteristics of the study cohort.

The median age in vaccinated group is 42.0 (95% CI 30.0, 56.0), and in unvaccinated group is 45.0 (95% CI 31.5, 58.5). 3 (23.1%) out of 14 unvaccinated individuals, but none of vaccinated individuals, developed severe COVID-19 disease. Sampling at 12 months after the third booster dose (T0), as well as at 2 weeks (T1) and 20 weeks (T2) after SARS-CoV-2 infection, allowed the immune trajectories to be measured to assess the establishment and maintenance of hybrid immunity. Simultaneous serum, peripheral blood mononuclear cell, and nasal swab samples were collected from each of the participants. The samples were subjected to a detailed analysis for both serological and cellular immune responses to SARS-CoV-2 antigens. The serum samples were tested for specific antibody titers using the enzyme-linked immunosorbent assay and neutralization activities against the ancestral, Delta, and Omicron BA.5 variants. We analyzed Fc-mediated phagocytosis activity to evaluate any functional differences in the antibody responses among vaccinated and unvaccinated individuals after BA.5 infection.

### Breakthrough infection enhanced the magnitude and durability of humoral antibody responses

Antibodies are considered as the correlates of protection against SARS-CoV-2 infection.^12^ Therefore, we investigated the magnitude of variant-specific immunoglobulin (Ig)G and IgA binding in the serum. First, the binding of IgG and IgA with ancestral, Delta, BA.5, XBB.1, and XBB.1.5 was assessed by analyzing the serum collected before infection and at 2 and 20 weeks after SARS-CoV-2 infection **(Figure 2a, 2b)**.

**Figure 2.**
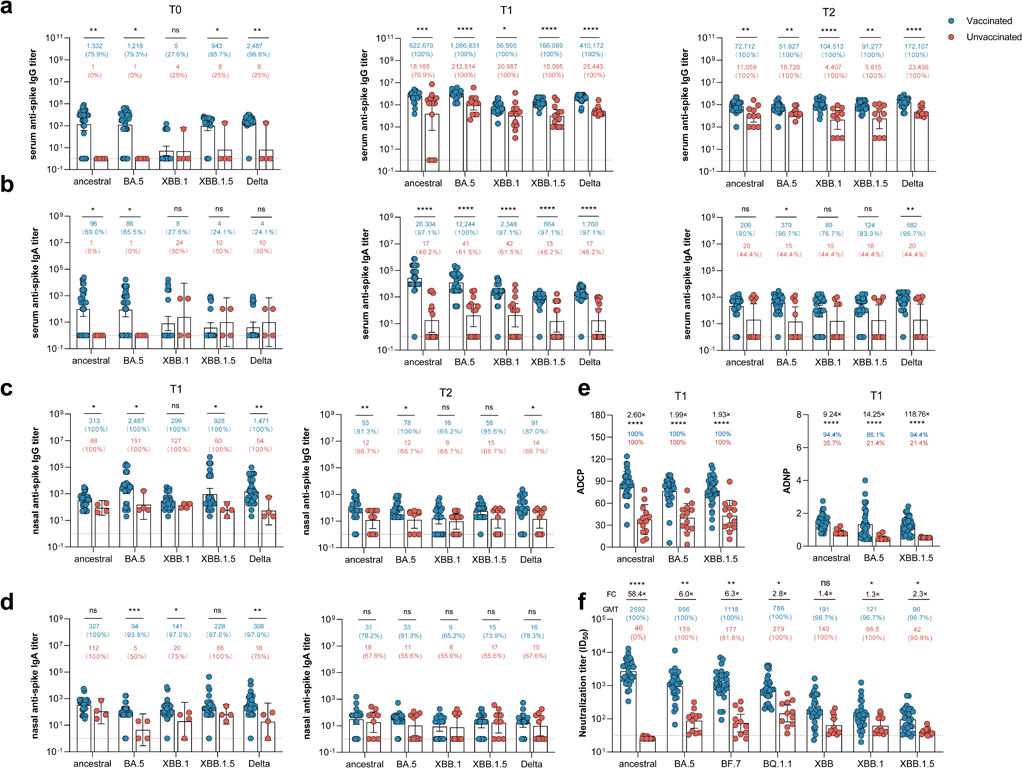
Omicron BA.5 infection enhanced serum and mucosal humoral response in triple CoronaVac vaccinated individuals. (**a-d**) Enzyme-linked immunosorbent assay (ELISA) measurement for serum anti-spike IgG titer (**a**), serum anti-spike IgA titer (**b**), nasal anti-spike IgG titer (**c**) and nasal anti-spike IgA titer (**d**) at different time points, including 12-month after the third dose vaccination (T0), 2 weeks (T1) and 20 weeks (T2) post BA.5 infection. (**e**) Antibody-dependent cellular phagocytosis (ADCP) and antibody-dependent neutrophil phagocytosis (ADNP) specific to spike protein of ancestral, BA.5 and XBB.1.5 at T1 were also analyzed. (**f**) Serum titers that achieved 50% pseudovirus neutralization (ID50) of seven SARS-CoV-2 variants including ancestral strain, BA.5, BF.7, BQ1.1, XBB, XBB.1, XBB.1.5, in vaccinated and unvaccinated individuals after BA.5 infection at T1 timepoint. Blue dots represent previous triple CoronaVac vaccinated individuals, and red dots represent unvaccinated individuals. Dotted lines indicated the lower limit of detection (LOD) for the assay. Data points on the bar graph represent individual titer and the line indicates geometric mean titer (GMT). GMTs of each group were noted on the top of each bar. Statistics were calculated using Mann-Whitney U test for single comparisons variables between groups. * indicates p <0.05, ** indicates p <0.01, *** indicates p <0.001, **** indicates p <0.0001; ns indicates no significant difference.

Baseline sera at T0 exhibited an ancestral spike-specific binding geometric median titer (GMT) of 1,332 (95% confidence interval [CI] 358–4,959), while the GMT specific to BA.5, XBB.1, and XBB.1.5 was 1,218 (95% CI 365–4,064), 5 (95% CI 2–13), and 2,134 (95% CI 1,295–3,519), respectively **(Figure 2a)**.

The convalescent serum from vaccinated individuals at T1 showed an increment of 672-fold for anti-ancestral spike IgG, with a GMT of 622,679 (95% CI 433,486– 894,445). The respective fold elevation in anti-BA.5, anti-XBB.1, and anti-XBB.1.5 spike IgG titers was 1164-fold (1,086,831, 95% CI 820,495–1,439,620), 9928-fold (56,595, 95% CI 39,632–80,819), and 234-fold (166,089, 95% CI 129,837–212,463), respectively. Compared to unvaccinated individuals, vaccinated individuals exhibited a considerably higher ancestral spike-specific IgG titer (622,679 vs. 15,127, p = 0.006) and BA.5 spike-specific IgG titer (1,046,394 vs. 212,514, p < 0.0001). Similar patterns were also observed for the XBB.1, XBB.1.5, and Delta spike-specific IgG titers.

We assessed the durability of the binding antibody titers at T2 (20 weeks after BA.5 infection). Compared with the unvaccinated controls, vaccinated individuals retained a significantly greater anti-spike IgG titer, not only specific to the ancestral spike (11,059 vs. 72,712, p = 0.004), but also to the BA.5 spike (18,726 vs. 51,827, p = 0.004), XBB.1 spike (4,407 vs. 10,4513, p < 0.0001), XBB.1.5 spike (5,815 vs. 91,277, p = 0.003), and Delta spike (23,456 vs. 172,107, p < 0.0001). Meanwhile, dynamic serological SARS-CoV-2-specific IgA responses were also detected (**Figure 2b**). After BA.5 infection, 40 of 41 fully vaccinated individuals (97%) were IgA-positive for the ancestral spike, and 100%, 97%, and 97% were IgA-positive for the BA.5, XBB.1, and XBB.1.5 spikes, respectively. 7 of 14 unvaccinated individuals (50%) were IgA-positive for the ancestral spike, and 62%, 62%, and 46% were IgA-positive for the BA.5, XBB.1, and XBB.1.5 spikes, respectively. Compared to T0, BA.5 infection induced strong recall responses in vaccinated individuals, leading to 1,547-fold, 299-fold, 56-fold, and 44-fold increased IgA titers specific to the ancestral, BA.5, XBB.1, and XBB.1.5 strains, respectively, at T1. Consistently, vaccinated individuals showed significant enhancement of spike-specific IgA responses across the tested variants compared with unvaccinated individuals (all p < 0.0001). Vaccinated individuals had significantly higher BA.5 spike and Delta spike IgA responses at T2 ((p=0.011 and p=0.009, respectively). Similar trends were also noted for the IgA titers specific to the ancestral, XBB.1, and XBB.1.5 spikes, but without significant differences. Collectively, our data suggest that a prior history of vaccination facilitates antibody recall, resulting in the boosted breadth and durability of SARS-CoV-2-specific IgG and IgA responses.

### Triple-dose CoronaVac expanded nasal IgA and IgG antibody titers after acquired BA.5 infection

Recent evidence has demonstrated that mucosal immunity plays a pivotal role in respiratory viral infection, and ancestral SARS-CoV-2 spike-specific mucosal IgA is protective against Omicron infection.^13^ To this end, we detected a higher nasal anti-spike IgA titer specific to BA.5 (94 vs. 5, p = 0.0005), Delta (308 vs.18, p = 0.005), and XBB.1 (141 vs. 20, p = 0.026) in the vaccinated group than in the unvaccinated group **(Figure 2c).** A similar trend was detected for nasal IgG responses, with substantially elevated the ancestral (313 vs. 88, p = 0.046), BA.5 (2487 vs. 151, p = 0.017), and XBB.1.5 (928 vs. 60, p = 0.014) IgA responses. However, such differences were less obvious at T2. Our data suggest that although most intramuscularly injected COVID-19 vaccines induced minimal anti-spike IgA responses, breakthrough infection via the mucosal route synergistically improved both systemic and mucosal anti-viral immunity, but the mucosal IgA response appeared to wane within 20 weeks.

### Breakthrough infection induced stronger antibody-dependent cellular and neutrophil phagocytosis than natural BA.5 infection

Fc-mediated effector functions have been demonstrated to reduce the severity of disease rather than reducing transmission; therefore, they might play a role in vaccine-attenuated disease.^14^ We recently reported that triple-dose CoronaVac could induce potent antibody-dependent cellular phagocytosis (ADCP) and antibody-dependent neutrophil phagocytosis (ADNP).^7^ The data show that the vaccinated group had substantially elevated ADCP and ADNP compared with unvaccinated controls **(Figure 2e)**. Specifically, plasma samples from triple-dose CoronaVac recipients showed higher ADCP responses specific to the ancestral (2.29-fold higher, p < 0.0001), BA.5 (1.91-fold higher, p < 0.0001), and XBB.1.5 (1.79-fold higher, p < 0.0001) spikes than unvaccinated controls. Minimal ADNP activity was detected among unvaccinated individuals, while the vaccinated individuals had considerably enhanced ancestral (1.75-fold higher, p < 0.0001), BA.5 (2.76-fold higher, p < 0.0001), and XBB.1.5 (2.44-fold higher, p < 0.0001) spike ADNP activity.

### Breakthrough infection enhanced the potency of neutralization activity against BA.5, XBB.1, and XBB.1.5

Omicron subvariants, especially XBB subvariants, are considered to be highly resistant to vaccines and infection-induced humoral immunity.^15,16^ Using the pseudovirus neutralization assay, the serum neutralization capability against the most tested variants was significantly increased in vaccinated individuals compared with unvaccinated controls. The geometric mean 50% neutralizing antibody titers (NT_50_ GMTs) for the ancestral, BA.5, BF.7, BQ1.1, XBB, XBB.1, and XBB.1.5 strains in the vaccinated group were 2,691.8, 955.8, 1,117.5, 785.9, 190.8, 121.4, and 95.9, respectively. The NT_50_ GMTs of the unvaccinated group against the ancestral (46.1; 58.4-fold, p < 0.0001), BA.5 (158.6; 6.0-fold, p = 0.002), BF.7 (177.4; 6.3-fold, p = 0.003), BQ1.1 (279.0; 2.8-fold, p = 0.032), XBB (140.1; 1.4-fold, p = 0.486), XBB.1 (96.5; 1.3-fold, p = 0.044), and XBB.1.5 (41.9; 2.3-fold, p = 0.023) strains were significantly compromised **(Figure 2f).** The largest difference was observed for neutralization activity against the ancestral strain, whereas the smallest difference was observed for the XBB.1 strain between vaccinated and unvaccinated individuals. The data suggest that BA.5 neutralization activity was effectively boosted once the virus had invaded the oropharynx among vaccinated individuals who received the ancestral strain-based CoronaVac. Further, immune escape of the XBB subvariants from Omicron convalescent serum was partly restored by immunization with triple-dose CoronaVac.

### BA.5 infection enhanced cross-reactive spike-specific B cell responses that persisted for up to 20 weeks

Beyond humoral immunity, cellular components are also particularly indispensable for long-term protection against COVID-19.^17^ We have shown previously that while antibodies decline over time, B cells persist and cross-react with SARS-CoV-2 variants of concern to some degree.^2,5^ Therefore, we evaluated the magnitude and cross-reactivity of the antigen-specific B cell response via flow cytometric numeration of B cells stained with differentially labeled ancestral, BA.1, BA.2, and BA.5 spike proteins **(Figure 3a)**. At baseline, ancestral and BA.5 spike reactive B cells were detectable in 23 (79%) and 17 (59%) of 29 participants, comprising a median of 0.090% (95% CI 0.049%–0.125%) and 0.047% (95% CI 0.024%–0.061%) of the total B cells, respectively, among vaccinated individuals. Minimal ancestral spike-specific B cells (0.030%, 95% CI 0.020%–0.036%) were detected in unvaccinated controls **(Figure 3a)**. Correspondingly, BA.5 spike-specific B cells increased from 0.047% of the total B cells at baseline to 0.43% at 2 weeks, and slightly decreased to 0.22% at 20 weeks after BA.5 infection. In contrast, BA.5 infected unvaccinated individuals had significantly lower levels of ancestral (1.67-fold lower, p = 0.018), BA.5 (1.79-fold lower, p = 0.002), and BA.1 (2.14-fold lower, p < 0.0001) spike-specific B cells at T1 compared with vaccinated individuals **(Figure 3b)**. Twenty weeks after BA.5 infection, unvaccinated individuals still exhibited a lower proportion of spike-reactive B cells than the vaccinated group across the tested Omicron subvariants, including BA.5 (1.91-fold lower, p = 0.007), BA.1 (2.19-fold lower, p < 0.0001), and BA.2 (1.95-fold lower, p = 0.014).

**Figure 3.**
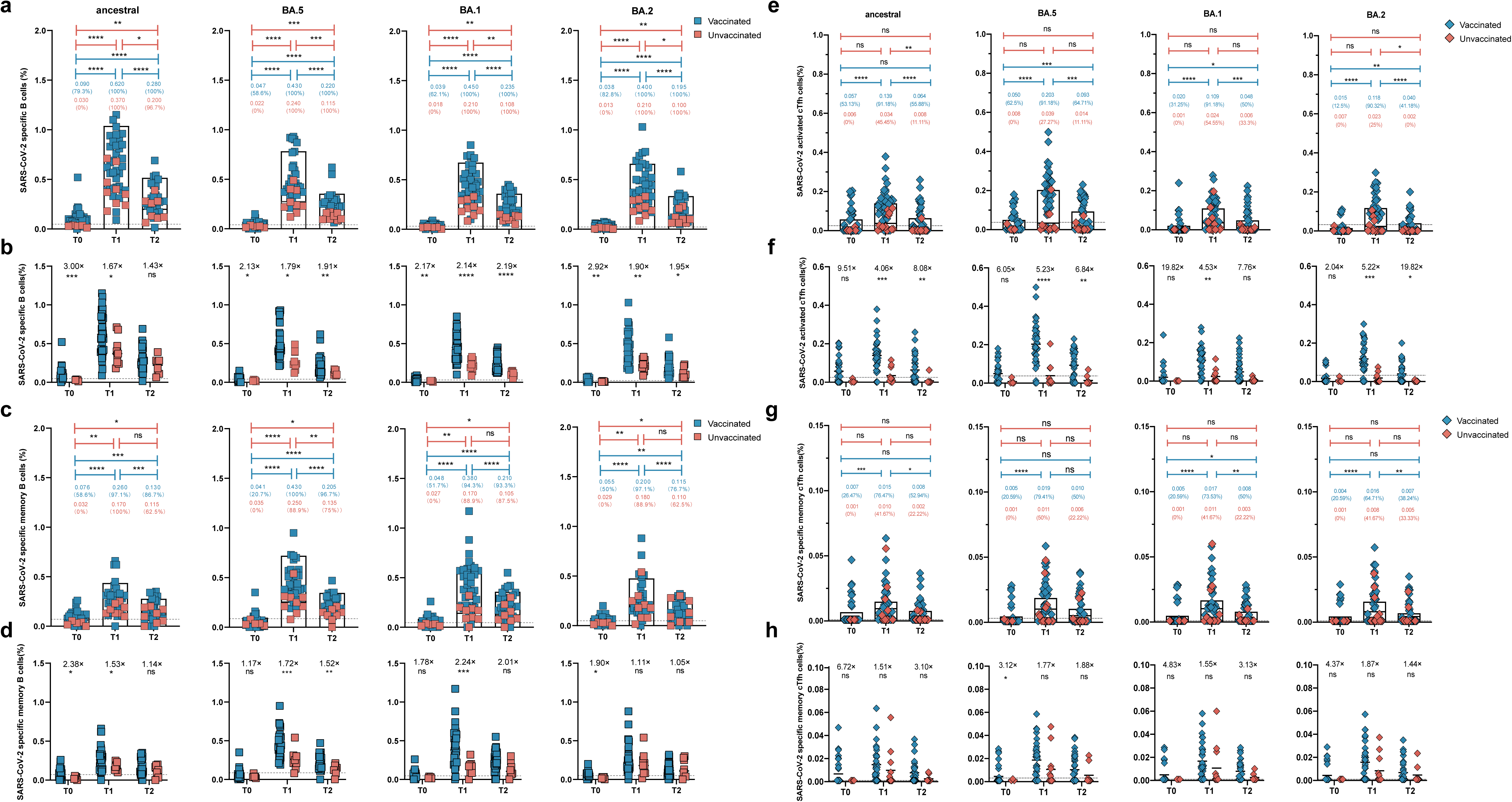
BA.5 infection enhanced cross-reactive spike specific B cell responses and T follicular helper (Tfh) cells that persist up to five months in vaccinated individuals. (**a, c, e, g**) The dynamic frequency of spike-specific B cells **(a)**, memory B cells **(c),** activated Tfh cells **(e)** and memory Tfh cells **(g)** specific to ancestral spike, BA.1 spike BA.2 spike and BA.5 spike at T0, T1 and T2 timepoint. Values above the symbols denote the median, and the percentage of positive responders were also noted. **(b, d, f, h)** The comparison of SARS-CoV-2 specific B cells **(b)**, memory B cells **(d),** activated Tfh cells **(f)** and memory Tfh cells **(h)** between vaccinated and unvaccinated individuals at T0, T1, and T2 timepoint. Fold change of spike-specific B cells between two groups at three timepoints. Data are shown as the fold-change between vaccinated versus unvaccinated donors. Dotted lines indicated the limit of detection (LOD) for the assay. Single comparisons variables between two groups were performed using the Mann-Whitney U test. Multiple comparisons of specific and specific memory B cell responses at three timepoints were performed using the Friedman’s one-way ANOVA with LSD. * indicates p <0.05, ** indicates p <0.01, *** indicates p<0.001, **** indicates p <0.0001; ns indicates no significant difference.

The proportion of spike-specific memory B cells among the total memory B cells was also analyzed **(Figure 3c).** Triple-dose CoronaVac recipients showed a median of 0.076%, 0.048%, 0.055%, and 0.041% memory B cells specific to ancestral, BA.1, BA.2, and BA.5, respectively, even at 12 months after the third booster vaccination. Memory B cells were absent in unvaccinated individuals at T0. BA.5 breakthrough infection expanded cross-reactive spike-specific memory B cells. At T1, compared to unvaccinated controls, vaccinated individuals showed remarkably higher levels of memory B cells specific to the ancestral (0.17% vs. 0.26%, p = 0.030), BA.5 (0.25% vs. 0.43%, p = 0.0009), BA.1 (0.17% vs. 0.38%, p = 0.0003), and BA.2 (0.18% vs. 0.20%, p = 0.451) strains **(Figure 3d)**. At T2, vaccinated individuals still maintained higher levels of spike-specific memory B cells than unvaccinated individuals, but the difference was not statistically different except for BA.5-specific memory B cells. Collectively, our data indicate that triple-dose CoronaVac established spike-specific memory B cells that could rapidly expand and proliferate into antibody-producing B cells upon breakthrough infection.

### Vaccination enabled activation of circulating follicular helper T (cTfh) cells and expansion of memory cTfh cells upon BA.5 breakthrough infection

Tfh cells play a vital role in the selection and activation of B cells into activated antibody-secreting cells at germinal centers (GCs), which is critical for the development of long-lasting, high-affinity antibody responses.^18^ A small population of cTfh cells resemble GC Tfh cells in terms of their phenotype and serve as a counterpart to GC Tfh cells to support antibody secretion.^18^ We tracked and depicted the frequency of spike-specific Tfh cells **(Figure 3e-h).** BA.5 infection expanded ancestral-specific cTfh cells from 0.057% (0.033%–0.081%) at baseline to 0.139% (0.110%–0.169%) at T1, and BA.5 spike-specific cTfh cells from 0.050% (0.030%–0.070%) at baseline to 0.203% (0.163%–0.242%) at T1, among vaccinated individuals **(Figure 3e)**. The frequency of spike-specific cTfh cells across SARS-CoV-2 variants was 2.04–19.82-fold higher in vaccinated individuals than in unvaccinated individuals. In particular, the percentage of BA.5-specific cTfh cells in the vaccinated group was 0.203% (0.163%–0.242%) and 0.093% (0.067%–0.120%) at T1 and T2, respectively (5.23-fold and 6.84-fold higher than in unvaccinated individuals) **(Figure 3f).**

At T0, despite being 12 months after the third dose of CoronaVac, vaccinated individuals not only showed a detectable level of ancestral spike-specific memory cTfh cell responses (0.007%, 95%CI 0.003%–0.011%), but they also showed a slightly lower frequency of memory cTfh cells that were cross-reactive to the BA.1, BA.2, and BA.5 spikes, ranging from 0.004% to 0.005% **(Figure 3g)**. There was a trend toward slightly higher spike-specific memory cTfh cells in vaccinated individuals than in unvaccinated individuals, but no significant difference was observed **(Figure 3h)**. Collectively, our data suggest that breakthrough infection could quickly recall and activate specific cTfh cells, but memory cTfh cells were not significantly elevated compared with unvaccinated individuals.

### BA.5 infection enhanced cross-reactive memory T cells among recipients of triple-dose CoronaVac

T cells contribute to the defense against viral infections by orchestrating antibody production and cytotoxic killing of infected cells.^19^ Previously, we observed that CoronaVac induced durable, cross-reactive T cell responses.^2,5^ Given their importance, we examined whether BA.5 breakthrough infection could quickly recall and expand ancestral- and BA.5-specific CD4^+^ and CD8^+^ T cell responses. The data show that triple-dose CoronaVac recipients still showed detectable CD4^+^ T cells with a median frequency of 0.026% (95% CI 0.016%–0.039%) **(Figure 4a)** and CD8^+^ T cells with a median frequency of 0.035% (95% CI 0.019%–0.056%) **(Figure 4b)**. In contrast, few spike-specific CD4^+^ or CD8^+^ T cell responses were detected in unvaccinated individuals. At T1, BA.5 breakthrough infection effectively expanded both CD4^+^ T cell (3.73-fold, p < 0.0001) and CD8^+^ T cell (5.45-fold, p < 0.0001) responses specific to the BA.5 spike **(Figure 4a, 4e).** Similar trends were also observed in the T cell responses across the ancestral strain and the Omicron subvariants at T2.

**Figure 4.**
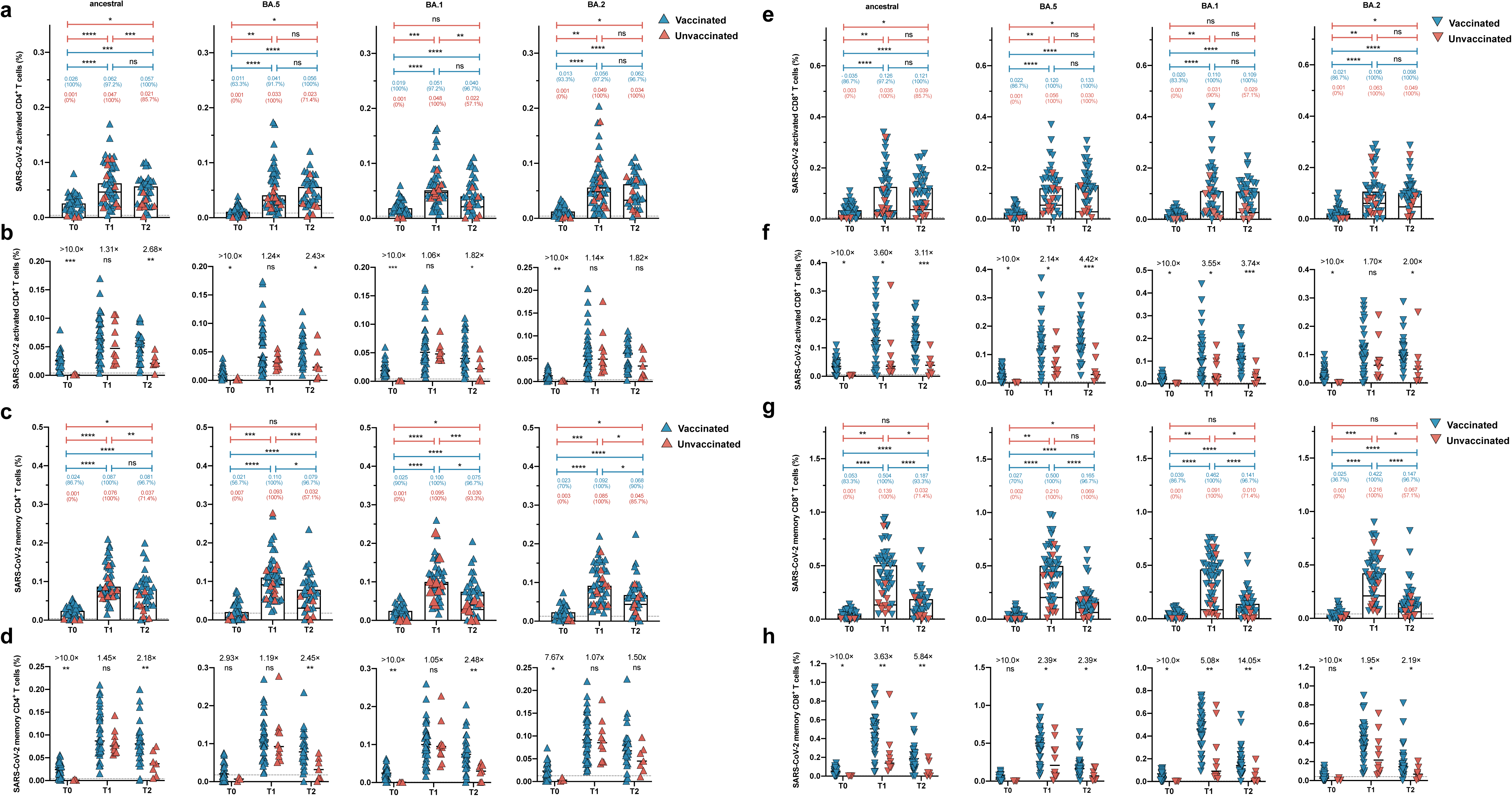
BA.5 infection enhanced cross-reactive memory T cells among previously triple CoronaVac vaccinees. (**a, c, e, g**) The dynamic change of activated CD4+ T cells **(a),** memory CD4+ T cells **(c),** activated CD8+ T cells **(e)** and memory CD8+ T cells **(g)** specific to ancestral, BA.1, BA.2 and BA.5 spike in vaccinated and unvaccinated groups at T0, T1, T2 timepoint. Values above the symbols denote the median and the percentage of positive responders were also noted. **(b, d, f, h)** The comparison of SARS-CoV-2 specific activated CD4+ T cells (b), memory CD4+ T cells (d), activated CD8+ T cells (f) and memory CD8+ T cells (h) between vaccinated and unvaccinated individuals at T0, T1, and T2 timepoint. Data are shown as the fold-change between vaccinated versus unvaccinated donors. Single comparisons variables between groups were performed using the Mann-Whitney U test. Multiple comparisons of specific T cell responses of three timpoints were performed using the Friedman’s one-way ANOVA with LSD. Dotted lines indicated the limit of detection (LOD) for the assay. Bars represented median value. p<0.05 was considered to be statistically significant. * indicates p < 0.05, ** indicates p < 0.01, *** indicates p < 0.001, **** indicates p < 0.0001. ns indicates no significant difference.

The memory T cell subpopulation is critical to determine anti-viral responses upon antigen exposure. Therefore, we longitudinally measured the frequency of specific memory T cell responses. We found that compared with the corresponding baseline samples, BA.5 breakthrough infection led to augmented BA.5 spike-specific memory CD4^+^ T cell (5.24-fold, p < 0.0001) and memory CD8^+^ T cell (18.52-fold, p < 0.0001) responses at T1 **(Figure 4c, 4g)**. Interestingly, there was a trend toward a slightly higher frequency of specific memory CD4^+^ T cell responses in vaccinated individuals than in unvaccinated controls, but without significant differences **(Figure 4d)**. In contrast, compared with unvaccinated controls, vaccinated individuals showed substantially higher levels of ancestral (0.504% vs. 0.139%, p = 0.001), BA.5 (0.500% vs. 0.210%, p = 0.011), BA.1 (0.462% vs. 0.091%, p = 0.002), and BA.2 (0.422% vs. 0.216%, p = 0.040) spike-specific CD8^+^ memory T cell responses **(Figure 4g, 4h).** To summarize, CD4^+^ and CD8^+^ memory T cells were still detectable 12 months after the third dose of CoronaVac. These pre-existing cross-reactive T cell responses and memory subsets could be effectively boosted and maintained by BA.5 infection.

### The increase in BA.5-specific immune responses after breakthrough infection was consistently higher than for the ancestral strain

Immune imprinting induced by ancestral strain-based vaccination might compromise the antibody response to Omicron-based boosters.^8,9^ Therefore, we compared the increases in the ancestral- and BA.5-specific immune responses in vaccinated individuals **(Figure 5)**. Notably, we found that the increase from T0 to T1 in the BA.5 spike-specific IgG response was substantially higher than for the ancestral spike **(Figure 5a)**. Similarly, the majority of BA.5-specific immune parameters, including B cells, memory B cells, memory Tfh cells, CD4^+^ T cells, memory CD4^+^ T cells, and memory CD8^+^ T cells, displayed substantially greater fold changes than those specific to the ancestral strain at T1 **(Figure 5b–6c).** Nevertheless, the enhanced increase in the BA.5-specific immune response was not observed at T2 **(Figure 5d–6f).**

**Figure 5.**
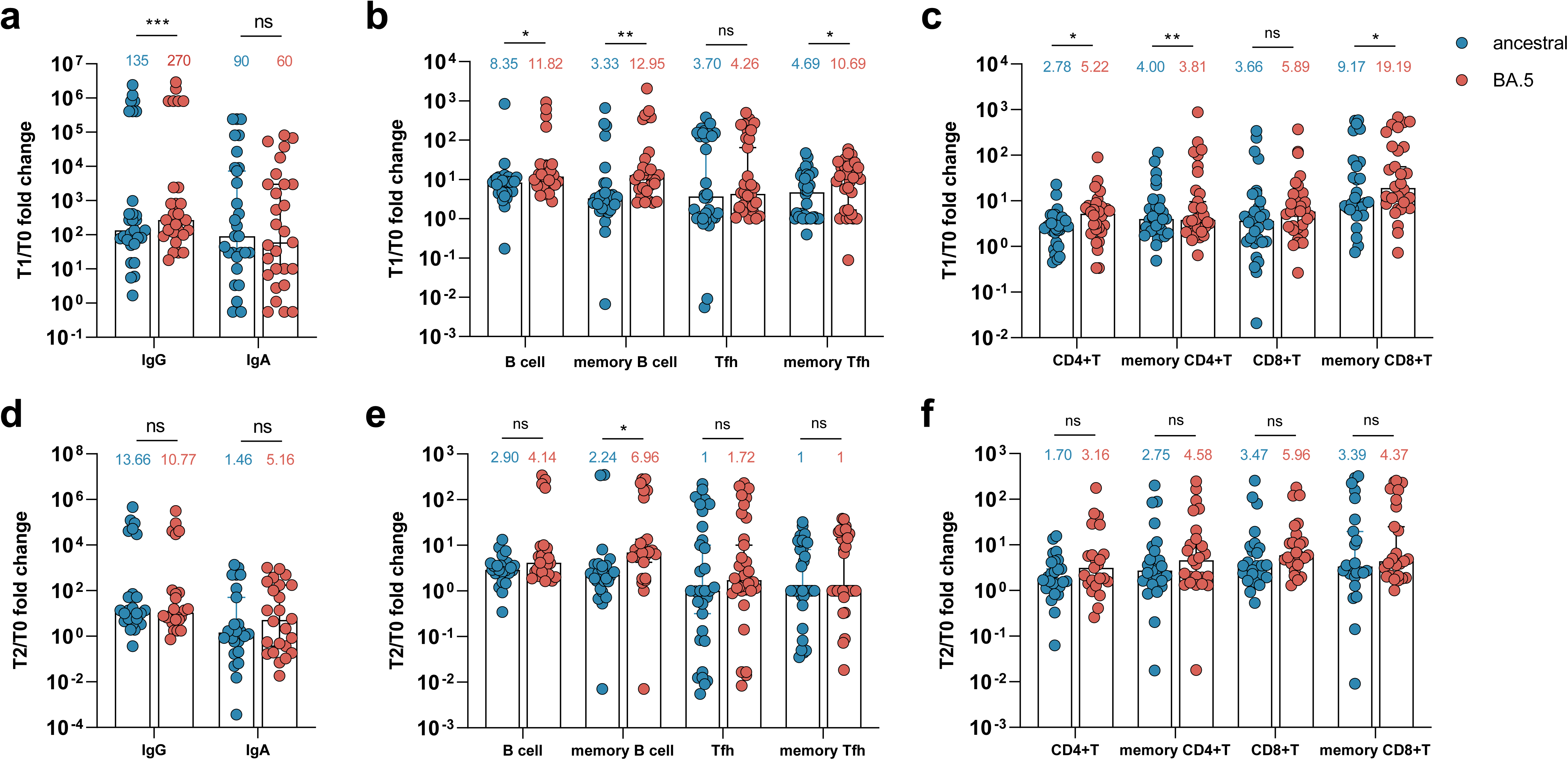
Increase fold of BA.5 specific immune responses post breakthrough infection were consistently higher than those of ancestral strain. (**a-c**) Fold change of of serum anti-spike IgG and IgA (a) and B cells, memoryB cells, Tfh and memory Tfh cells (b), CD4+T cells, memory CD4+T cells, CD8+T cells and memory CD8+T cells (c) among vaccinated group against ancestral and BA.5 strain from T1 timepoint versus T0 timepoint. (**d-f**) Fold change of of serum anti-spike IgG and IgA (a) and B cells, memoryB cells, Tfh and memory Tfh cells (b), CD4+T cells, memory CD4+T cells, CD8+T cells and memory CD8+T cells (c) among vaccinated group against ancestral and BA.5 strain from T2 timepoint versus T0 timepoint. Fold change of of serum anti-spike IgG and IgA (a) and B cells, memoryB cells, Tfh and memory Tfh cells (b), CD4+T cells, memory CD4+T cells, CD8+T cells and memory CD8+T cells (c) among vaccinated group against ancestral and BA.5 strain from T1 timepoint versus T0 timepoint. Data are shown as the fold-change between T0, T1, and T2 in vaccinated group. Wilcoxon matched-pairs signed rank test was performed for paired data. * indicates p < 0.05, ** indicates p < 0.01, *** indicates p < 0.001, **** indicates p < 0.0001. ns indicates no significant difference.

**Figure 6.**
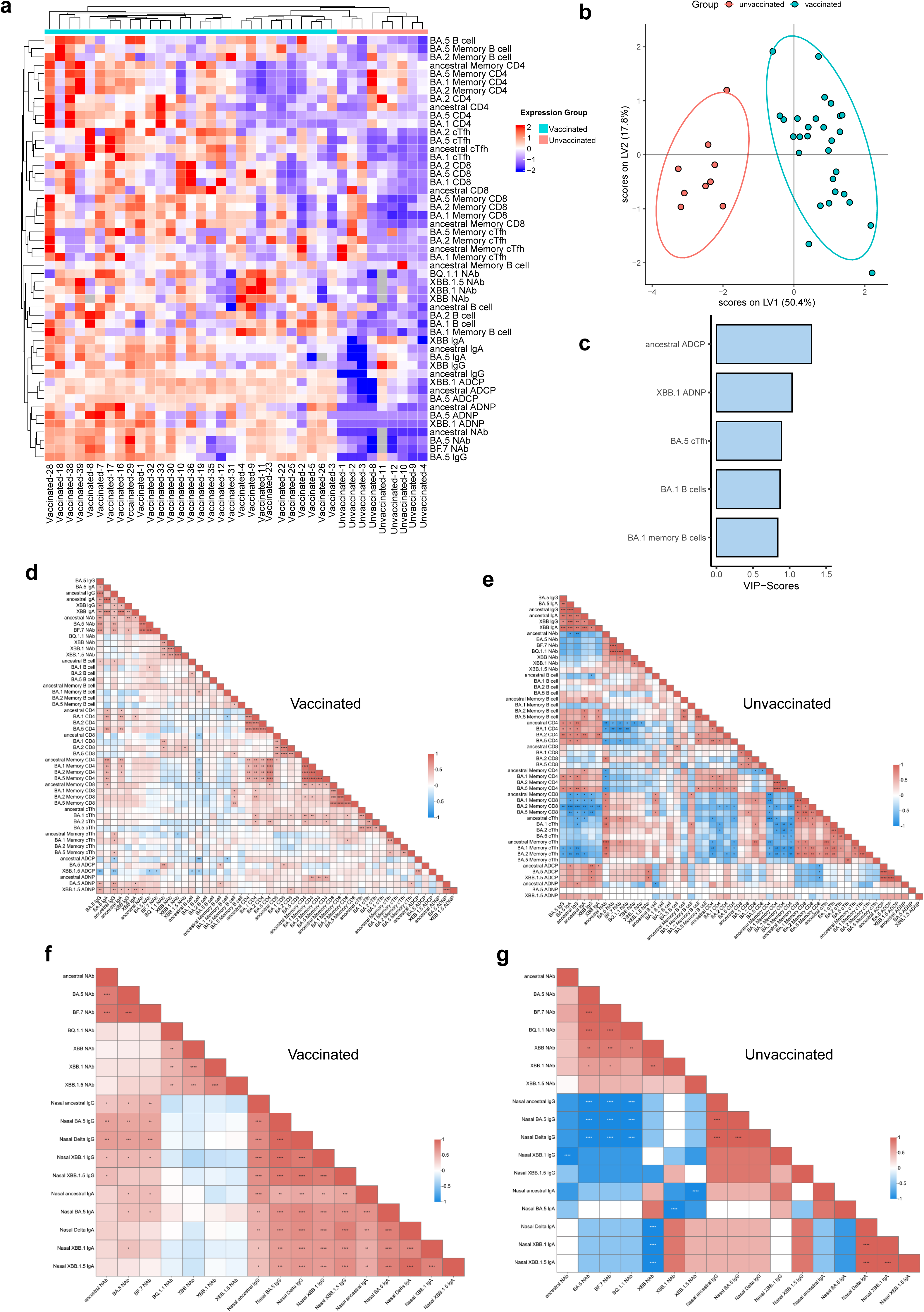
Distinct patterns of immune trajectories in breakthrough infected individuals versus natural infected donors. (**a**) A cold-to-hot Heatmap and hierarchical clustering represents the scaled magnitude of SARS-CoV-2 specific humoral, mucosal and cellular immune responses including spike binding antibody, neutralization activities, Fc-mediated effector function, cellular responses, at T1 timepoint after BA.5 infection between vaccinated versus unvaccinated group. Each column represents a vaccinated or unvaccinated individuals, while each role represents an immune parameter. The distribution of markers and participants in the cohort was automatically performed by supervised hierarchical clustering. **(b)** A Least Absolute Shrinkage Selection Operator (LASSO) was used to select antibody features that contributed most of the discriminate vaccinated versus unvaccinated individuals. A partial least square discriminant analysis (PLSDA) was used to visualize samples. LASSO selected features were ranked based on their Variable of Importance (VIP) score, and the loadings of the latent variable 1 (LV1) were visualized in a bar graph. **(c)** Selected immune parameter features were ordered according to their variable importance in projection (VIP) score. (**d-e**) Correlation analysis between the parameters of adaptive immune responses, including binding antibody, neutralization activities, B cell, CD4 cell, CD8 cell, Tfh cell responses as well as their memory subsets at T1 timepoint after BA.5 infection in vaccinated **(d)** versus unvaccinated donors **(e).** (**f-g**) Correlation analysis of mucosal antibody response and serum neutralization activity in vaccinated individuals **(f)** versus unvaccinated **(g)** donors. The strength of a correlation (Sperman’s correlation coefficient) is demonstrated by color of square, the significance is indicated by asterisks. Correlation between 2 continuous variables was analyzed using the spearman correlation analysis. * indicates p < 0.05, ** indicates p < 0.01, *** indicates p < 0.001, **** indicates p < 0.0001. ns indicates no significant difference.

### Distinct immune trajectories in breakthrough-infected individuals versus naturally infected individuals

The BA.5 infected vaccinated group exhibited remarkable divergence in SARS-CoV-2-specific immune responses compared with the unvaccinated group. Therefore, further exploratory analyses were performed to gain a more detailed understanding of their fundamental differences. Following centering and scaling, hierarchical clustering of the participants revealed that vaccinated samples clustered away from the unvaccinated samples (**Figure 6a)**, highlighting the potent multicomponent immune memory induced by CoronaVac, even up to 12 months after the third CoronaVac dose. A multivariate partial least squares-discriminant analysis was performed across the vaccinated group and the unvaccianted group at the peak immune response after breakthrough infection **(Figure 6b).** We found that vaccinated individuals could be perfectly separated from unvaccinated individuals. Notably, the immune profiles of participants in the vaccinated group exhibited selective enrichment of ancestral-specific ADCP, XBB.1-specific ADNP, BA.5-specific Tfh cells, and BA.1-specific B cells and memory B cells **(Figure 6c).**

We also performed correlation analyses for the measured immune parameters, including antibody binding, neutralization activity, B cells, CD4^+^ cells, CD8^+^ cells, and Tfh cell responses, as well as their memory subsets, among the vaccinated group and the unvaccinated group. In the vaccinated group, we observed a moderate-to-low degree of correlation among the parameters defining the antibody response and among the parameters defining T cell responses. Further, binding antibody responses were positively correlated with the CD4^+^ T cell response and the corresponding memory subsets **(Figure 6d)**. In the unvaccinated group, we observed a strong or moderate degree of correlation within the antibody compartment and within the T cell compartment. Nevertheless, a considerable proportion of inverse correlations were identified between antibodies and CD8^+^ memory T cells and cTfh cells, as well as between the CD4^+^ T cell response and memory Tfh cells (**Figure 6e**). These results suggest that breakthrough infection had a well-integrated adaptive response between the antibody compartment and the T cell compartment, whereas the unvaccinated group harbored less coordinated immune trajectories between the antibody and T cell compartments, which might contribute to suboptimal antibody responses.

Additionally, we investigated the correlation between systemic and mucosal humoral immunity. Interestingly, BA.5-infected individuals in the vaccinated group demonstrated a significant positive correlation between serum-neutralizing activity and nasal IgA binding as well as IgG responses. Further, strong positive correlations were identified within nasal IgA and IgG binding specific to SARS-CoV-2 variant spike antigens **(Figure 6f)**. Surprisingly, for previously unvaccinated individuals, serum neutralizing activity was negatively correlated with nasal binding antibody titers after BA.5 infection. Additionally, few cross-correlations were identified among the parameters of the nasal IgG and IgA binding antibodies **(Figure 6g).** Collectively, our data highlight that vaccinated individuals have better synchronization of multiple immune components than unvaccinated individuals upon heterologous SARS-CoV-2 infection.

## Discussion

The inactivated whole-virion vaccine CoronaVac is one of the most widely used COVID-19 vaccines worldwide.^20^ Although there has been continual evolution of viral variants, which have managed to evade antibody responses to varying degrees, triple-dose CoronaVac has retained more effectiveness against severe disease than against overall infection.^21^ Emerging evidence suggests that both T cells and antibody responses provide the greatest protection against infection and death in severe COVID-19 cases.^22^ In this study, we provide strong evidence of the robust immune memory responses to ancestral and Omicron subvariants after triple-dose CoronaVac, which are maintained 12 months after the last dose. We also performed head-to-head comparisons of immune responses among vaccinated individuals versus unvaccinated individuals. When considered alongside the risk of severe infection and the long-term consequences of infection, our findings have important implications for vaccine policy.

Understanding the durability of the protection conferred by SARS-CoV-2 vaccination is of utmost importance to guide COVID-19 mitigation policies worldwide. The first key question we addressed is the durability of the immune memory and whether it is likely to protect against severe COVID-19 in the long term. We showed that serum anti-spike IgG and IgA responses were still detectable among the majority of triple-dose CoronaVac recipients at 12 months after the last dose. Additionally, we found that B and T cell responses after triple-dose CoronaVac were durable up to 12 months after the third dose, and that ancestral and Omicron subvariants were equally well recognized. Taken together, our longitudinal follow-up over 12 months after the third CoronaVac dose demonstrated the durability and long-term maintenance of SARS-CoV-2 circulating B cells and CD4^+^ and CD8^+^ memory T cells.

The second key question that we attempted to address is the impact of immune imprinting from triple-dose CoronaVac on the adaptive response against Omicron subvariants. Our study analyzed breakthrough infection-induced humoral responses and cellular responses simultaneously, which allowed us to profile the dynamic kinetics of the antibody and cellular compartments. As the majority of people are vaccinated against the ancestral SARS-CoV-2 strain, immune imprinting induced by ancestral strain-based vaccination presents a major challenge to the performance of updated boosters. It has been suggested that boosting with a variant that is antigenically distinct from the ancestral strain would recall pre-existing memory B cells induced by the ancestral strain, preventing the generation of humoral immune responses targeting new variants.^23^ Intriguingly, our results did not show strong evidence of antigenic sin in the antibody responses after Omicron infection. We speculate that this might be due to the relatively longer interval between the last booster dose and breakthrough infection than in previous reports. However, this warrants further investigation.^9,24^ Additionally, we showed that the increases in BA.5-specific memory cellular responses, including memory B cells, Tfh cells, CD4^+^ T cells, and CD8^+^ T cells, after breakthrough infection were consistently higher than those of the ancestral strain. This finding aligns with recent evidence showing that either SARS-CoV-2 Omicron boosting or Omicron breakthrough infection not only triggers rapid memory cellular responses but also induces *de novo* B^25^ and T cell^26^ responses. In addition, we^2,5^ and others^28^ have shown that T cell responses are less affected by viral variants than antibodies, which is likely due to the broader range of epitopes available to T cells compared with antibodies, where protective responses are more focused.

Further support for the protective role of triple-dose CoronaVac comes from our observation that hybrid immunity strengthens and broadens antibody responses, mucosal immunity, and cellular immunity, providing the most sustained protection. Despite there being a large amount of evidence highlighting the remarkable enhancement of serological and cellular responses elicited by hybrid immunity, there is a paucity of data suggesting that mucosal immunity could also be strengthened. For the first time, our data demonstrate that a larger proportion of breakthrough-infected individuals harbored respiratory spike-specific IgA responses than naturally infected individuals. Additionally, spike-specific nasal IgG and IgA responses of remarkable magnitude were also identified in individuals with hybrid immunity. The augmented mucosal immunity, especially the nasal IgA response,^28^ might be strongly associated with superior protection against SARS-CoV-2 reinfection, as described previously.^29^

Our study has important implications for the vulnerability of unvaccinated Omicron-infected individuals to reinfection by circulating and emerging SARS-CoV-2. In the absence of vaccination, BA.5 infection-elicited humoral responses showed remarkably reduced serum neutralization activity against BA.5 and BF.7, as well as a subtle decrease against BQ1.1, XBB, XBB.1, and XBB.1.5, and dramatic loss against the ancestral strain. Additionally, despite infection via the respiratory route, the BA.5 spike nasal IgA and IgG titers were dramatically lower in unvaccinated controls than in vaccinated individuals after BA.5 infection, indicating the possible risk of reinfection. We extended our analysis to Fc-mediated phagocytosis and mucosal immunity. In particular, we found that BA.5-specific ADNP responses and nasal IgA responses were not effectively induced in unvaccinated individuals after BA.5 infection. In contrast, the BA.5-specific T cell response was only subtly lower in unvaccinated individuals than in vaccinated individuals. Taken together, the modest immune responses in BA.5-infected unvaccinated individuals leave this unvaccinated group at risk of being reinfected with Omicron subvariants. Our data indicate that Omicron-based vaccines might not be an ideal immunogen in SARS-CoV-2-naïve individuals.

For the first time, our results provide an immune landscape of hybrid immunity elicited by BA.5 infection in triple-vaccinated versus unvaccinated individuals. We showed that vaccination with triple-dose CoronaVac effectively evoked higher-quality immune responses characterized by convergent development of cross-reactive humoral and cellular immune compartments, as well as collaborative systemic and mucosal antibody compartments. Machine learning analysis showed that vaccinated individuals generated potent immune responses involving ADCP, ADNP, Tfh cells, activated B cells, and memory B cells, with distinct pattern from unvaccinated donors. We also suggest that a one-time Omicron BA.5 infection might not be sufficient to trigger cross-reactive humoral and cellular responses.

This study has some limitations that should be noted. First, T and B cells in the peripheral blood only account for a small proportion of the T and B cell population in the body. Therefore, the measurement of circulating cellular responses might not fully represent the landscape of T cell immunity in vaccinated individuals, especially tissue-resident memory T cells. Second, the participants in our cohort were generally young; older individuals were not included. Third, we used peptide stimulation to quantitatively measure T and B cell responses to SARS-CoV-2 spike proteins by flow cytometry. The use of conjugated peptide major histocompatibility complex (pMHC) multimers might result in the sensitive detection of antigen-specific T cells at a higher resolution. Finally, analysis of a more diverse cohort might facilitate further dissection of the immune responses induced by other types of vaccination.

In summary, we have shown that triple-dose CoronaVac remarkably augments antibody, mucosal, and cellular responses that are potent, durable, and cross-reactive to Omicron subvariants. This study sheds light on the dynamics of human adaptive immunity to SARS-CoV-2 Omicron subvariants in a highly vaccinated population with inactivated COVID-19 vaccines.

## Materials and Methods

### Study design

The purpose of this prospective, observational study (NCT05680896) was to directly compare the humoral and cellular immune response among triple CoronaVac vaccinated individuals or unvaccinated individuals who were subsequently contracted Omicron BA.5 infection during the omicron wave last Dec in China. Serum samples were collected from participants, which were analyzed using enzyme-linked immunosorbent (ELISA) assay, pseudovirus neutralization assay, antibody dependent cellular phagocytosis, as well as flow-cytometry based cellular analysis.

### Cohort selection and sample collection

Healthcare workers at Nanjing Drum Tower Hospital were recruited and enrolled in the study belonging to two groups: vaccinated and unvaccinated individuals. Written informed consent was obtained at the time of enrollment and study approval was obtained from the ethics committee of institutional review board from Nanjing Drum Tower Hospital (2021-034-01 and 20222-746). The vaccinated group received the first two doses of CoronaVac in Feb 2021 and the third dose of CoronaVac in Nov 2022, while the unvaccinated individuals did not receive any COVID-19 vaccine prior to Omicron BA.5 infection. Both groups had PCR confirmed diagnosis of COVID-19 during the recent omicron wave from Dec 2022 and Jan 2023 and the infected viral sequence was further confirmed by next-generation sequencing. For vaccinated participants, baseline samples were collected at 12-months post the third CoronaVac immunization. The post infection samples were collected at the median of 18 days (15-21) and 146.5 days (144.8-150.0) post BA.5 infection in vaccinated group, and 14 days (13.5-15) and 127.5 days (113.8-140.3) in unvaccinated group, respectively.

Sera were obtained by collecting 4-6 mL of whole blood in a BD Vacutainer Plus Plastic Serum Tube, which was centrifuged for 10 min at 1000xg before serum was aliquoted and stored at −20°C. Meanwhile, the nasal swabs were also collected for measurement of specific mucosal antibody titers. For the cellular analysis, PBMC were isolated from blood collected in ethylenediaminetetraacetic acid-anticoagulated (EDTA) tubes by lymphocyte separation medium density gradients (Stemcell technologies, cat# 07801) and resuspended in PRMI 1640 medium supplemented with 10% FCS, 1% penicillin/streptomycin and 1.5% HEPES buffer (complete medium; Thermo scientific) for stimulation assays or stored at - 135°C until used.

### Protein and peptides

Pools of 15-mer peptides overlapping by 11 amino acid (aa) and together spanning the entire sequence of SARS-CoV-2 Spike glycoprotein (S) from ancestral, Omicron BA.1 (B.1.1.529), BA.2 (B.1.1.529.2) and BA.5 (B.1.1.529.5) variants for *ex vivo* stimulation of PBMC for flow cytometry. The ectodomain of ancestral SARS-CoV-2 spike (GenBank: MN908947.3) was expressed as previously described.^10^ The protein was purified from FreeStyle 293-F cells (Thermo Fisher Scientific, USA) using affinity chromatography followed by size exclusion chromatography, detailed as described previously.^10^

### Measurement of serum and nasal SARS-CoV-2 spike-specific humoral responses

Serum or nasal swabs for immunological assessments were taken at three different time points, including 12 months post the third dose before BA.5 infection, 2 weeks and 20 weeks post BA.5 infection.^2,10^ Antigen-specific serological and nasal swab antibodies against SARS-CoV-2 were determined by an in-house enzyme-linked immunosorbent assay (ELISA), as previously described. Antibody-dependent cellular phagocytosis (ADCP) and antibody-dependent neutrophil phagocytosis (ADNP) were described in the previously study.^7^ Pseudovirus neutralization assay was performed using lentivirus-based SARS-CoV-2 pseudoviruses were provided by Vazyme Biotech Co., Ltd, as previously described.^2,11^

### Antigen-specific Measurement of Cellular Analyses

The biotinylated ectodomain of spike protein was fluorescently labeled to identify SARS-CoV-2 specific circulating B cells and memory B cells. The detailed approach has been described in previous studies.^2,5^ To measure antigen specific circulating and memory CD4^+^, CD8^+^ T cells and Tfh cells, activation-induced marker (AIM) assay were performed as previous described.^2,5^ Stained samples were acquired on a fluoresence-activated cell sorter (FACS) FACSAria™ III Cell Sorter instrument (BD Biosciences) and analyzed using FlowJo software version 10.7.1 (FlowJo LLC, BD Bioscience).

### Data analysis

To show the potential distinction of immune parameters between two groups, a cold-to-hot hierarchical clustering heatmap represents the scaled magnitude of SARS-CoV-2 specific humoral, mucosal and cellular immune responses including spike binding antibody titers, neutralization activities, Fc-mediated effector function, cellular responses, at T1 timepoint between vaccinated versus unvaccinated group. Each column represents a vaccinated or unvaccinated individual, while each row represents an immune parameter. The distribution of markers and participants in the cohort was automatically performed by supervised hierarchical clustering. Complexheatmap R package (2.15.1) was used for analysis.

A multivariate classification model was built to discriminate immunological profiles among triple vaccinated and unvaccinated groups using tested adaptive immune parameters in our study. All data were normalized using z-scoring before analysis. Feature selection was performed using least absolute shrinkage and selection operator (LASSO). Partial least square discriminant analysis (PLS-DA) was used for classification and visualization of immune parameters from two groups. Selected immune features were ordered based on Variable Importance in Projection (VIP) score and the first two latent variables (LVs) of the PLS-DA model were used to visualization. R package ‘ropls’ version 1.20.043 and ‘‘glmnet’’ version 4.0.244 and the systemseRology R package (v.1.1) (https://github.com/LoosC/systemsseRology) were used for analysis.

For the cross-correlation analysis, spearman correlation analysis was used for correlation analysis between all tested immune parameters. The correlation analysis was presented using ChiPlot (https://www.chiplot.online/correlation_heatmap.html).

### Statistical analysis

Binding antibody titers or neutralizing titers were expressed as geometric mean titers (GMTs). The mean (standard deviation) or median (95% confidence interval (CI)) was used to present the continuous variables. Categorical variables were described as the counts and percentages. Single comparison variables between groups were performed using the Mann-Whitney U test. Multiple comparisons of antibody titers, specific activated or memory B cell and T cell responses were performed using the Friedman’s one-way ANOVA with LSD. Correlation between 2 continuous variables was analyzed using the Spearman correlation analysis. p<0.05 was considered to be statistically significant.* indicates p < 0.05, ** indicates p < 0.01, *** indicates p < 0.001, **** indicates p < 0.0001. ns indicates no significant difference. SPSS software program version 22.0 (Chicago, IL, USA) were used to for data analysis.

## Data availability

R package ‘ropls’ version 1.20.043 and ‘‘glmnet’’ version 4.0.244 and the systemseRology R package (v.1.1) (https://github.com/LoosC/systemsseRology) were used for analysis.

The correlation analysis was presented using ChiPlot (https://www.chiplot.online/correlation_heatmap.html).

## Author contributions

Y.C., C.W. and M.N.contributed to study design, consultation and funding. M.N., Y.C., T.Z. and C.L. contributed to study design, experiments, manuscript writing and revision; Y.G, L.C., W.L. and S.Y. contributed to experiments, data analysis, bioinformatics analysis and figure generation; S.Y. and Y.T. contributed to experiments; J.N. and M.Z. contributed to bioinformatics analysis; Q.L., G.Z. and G.J. contributed to critical technical supports and reagents; M.Z. and W.L. contributed to consultation; All authors have read and approved the article.

## Acknowledgments

We are grateful to all the participants who participants in this study.

